# Differential myoblast and tenoblast affinity to collagen, fibrin and mixed threads in the prospect of muscle-tendon junction modelisation

**DOI:** 10.1101/2020.05.12.091868

**Authors:** Clément Rieu, Nicolas Rose, Anissa Taleb, Gervaise Mosser, Bernard Haye, Thibaud Coradin, Fabien Le Grand, Léa Trichet

## Abstract

The myotendinous junction transfers forces from muscle to tendon. As such, it must hold two tissues of completely different biological and cellular compositions as well as mechanical properties (kPa-MPa to MPa-GPa) and is subject to frequent stresses of high amplitude. This region remains a weak point of the muscle-tendon unit and is involved in frequent injuries. We here produce fibrin (40 mg/mL, E0 =0.10 ± 0.02 MPa) and collagen (60 mg/mL, E0=0.57 ± 0.05 MPa) threads as well as mixed collagen:fibrin threads (3:2 in mass, E0 = 0.33 ± 0.05 MPa) and investigate the difference of affinity between primary murine myoblasts and tenoblasts. We demonstrate a similar behavior of cells on mixed and fibrin threads with high adherence of tenoblasts and myoblasts, in comparison to collagen threads that promote high adherence and proliferation of tenoblasts but not of myoblasts. Besides, we show that myoblasts on threads differentiate but do not fuse, on the contrary to 2D control substrates, raising the question of the effect of substrate curvature on the ability of myoblasts to fuse *in vitro*.

## Introduction

Tendons are connective tissues that anchor skeletal muscles to the bones, transmitting the force generated by the muscles to the skeletal structure. As such, tendons play an obvious role in everyday movement and posture control. In case of injury, partly due to their low vascularization, they exhibit poor regeneration capacity resulting in loss of mobility, pain, *etc.* Injury of the muscle-tendon unit often occurs in the junction area as reported in many articles ^1–5^. In particular, muscle injuries are often located close to the junction ^6,7^, leading to the rupture of the myofibres rather than the tearing of the collagen fibres from the sarcolemma. Tensile tests on whole muscle-tendon units from rabbits also highlighted the weakness of the myo-tendinous junction (MTJ) area ^6^. As muscles often taper in contact with the tendon, stress concentration increases with decreasing cross-section ^8^.

During muscle regeneration process, quiescent muscle stem cells called satellite cells activate, and give rise to proliferating myoblasts destined to differentiate into myocytes able to fuse with one another or with injured myofibres ^9^. Up to date both the intrinsic transcription factors and the extrinsic cell-to-cell signaling pathways controlling muscle stem cells and myoblasts functions are well characterized. However, an emerging body of evidence supports the notion that myogenic cells integrate biochemical factors together with biomechanical cues that may control their cell fate^10^.

Muscles and tendons have completely different composition and structure, resulting in radically different mechanical properties: muscles are soft materials with Young moduli between 0.005 and 3 MPa while tendons are much stiffer, with Young moduli ranging from 500 to 1850 MPa ^11^. Junctions between materials with such a difference in mechanical properties may result in high interfacial stress and are likely to fail first during mechanical solicitation all the more since the stress borne by the MTJ reaches several tens of kPa ^12^. The increase of the contact surface between tendon and muscle by a factor 10 to 20, with finger-like structures, enables to decrease this stress ^13^. Along the myofibers the repeating sarcomeric units consist of an overlap of myosin and actin filaments, with Z-lines acting as anchoring points of the actin filaments and delineating the sarcomeric segments. In the peripheral finger-like structures, the last Z line is linked to the interdigitated sarcolemma through actin filament bundles. These actin filaments bind to transmembrane protein complexes containing integrins, and interact with the collagen IV and laminin-rich basal membrane and the extra-cellular matrix of tendons, rich in collagen I. Rebuilding the MTJ following muscle injury may thus subject progenitor cells to highly different environmental stiffnesses that promote divergent cellular responses.

For both muscular and tendon sides, reconstruction of the MTJ is relevant to help reconstruction in case of damaged MTJ or for better integration of the neo-tissue. For instance, in the hypothetical case of the grafting of a tendon *de novo*, integration at the muscular level through a new MTJ may be more effective and rapid as muscle is highly vascularized and is rich in satellite cells. MTJ structure is much less documented than muscle or tendon ones, and attempts to reproduce the MTJ are also scarce. We can cite the work reported by Ladd *et al.* ^11^ in which two different polymers are electrospun with collagen the poly(L lactic acid), being stiffer, for the tendon part and poly(ε -caprolactone) for the muscle part. Merceron *et al.* ^14^ used co-printing of two synthetic polymers polyurethane (PU) and poly(ε-caprolactone) (PCL) together with a bio-ink based on hyaluronic acid, fibrinogen and gelatin. The bio-ink contained C2C12 myoblasts on the PU side while the bio-ink on the stiffer PCL side contained NIH/3T3 cells. Both works used two materials with different mechanical properties to mimic the tendon and the muscle sides.

Mechanical properties of the substrate have been highlighted as key factors to control cell migration ^15,16^ and differentiation ^17,18^. In particular, substrate stiffness has been shown to play a role in muscle stem cells self-renewal ^19^, as well as in myotubes differentiation ^20^. Substrate stiffness has also been shown to promote tendon progenitor cells (here called tenoblasts) differentiation ^21,22^. Besides cell interactions with structural proteins have been shown to influence cell phenotype ^23,24^. Among them, collagen and fibrin are among the preferential proteins used in tissue engineering.

Type I collagen is an ubiquitous fibrillar protein found in most of the connective tissues *in vivo* ^25^. It is widely used in biomaterials ^26^ as it is a relatively stiff body protein ^27^ that bears multiple adhesion sites ^28,29^ for integrin receptors ^30^. In particular, it is the major component of tendons, reaching 55 to 80 % of tendon’s dry weight ^31^. On the other hand, fibrin is a major player in haemostasis, and polymerizes to form a clot after activation of its precursor, the blood-born fibrinogen, by thrombin ^32^. This fibrillar protein exhibits high deformability ^33^ and direct integrin adhesion sites ^34–36^, making it a material of choice for various applications such as for cardiovascular remediation ^37–39^ or cartilage repair ^40–42^. It is particularly favored for muscular reconstruction ^43–45^.

In this work, we investigate the effect of collagen I, fibrin and of mixed collagen:fibrin materials on mouse primary tenoblasts and myoblasts. Given the differing mechanical properties of the constitutive proteins, such scaffolds, composed only of biopolymers, might represent new types of tools to screen the respective influence of compositional and mechanical features on cell behavior. As a first step and in the prospect of musculotendinous junction modelisation, the capacity of highly concentrated collagen to offer a proper environment for tenoblasts, and of softer fibrin to favour muscular development, will be tested, as well as the influence of a mixed material. For this purpose, collagen, fibrin and “mix” threads are extruded. Mechanical properties and structure of threads are characterized and primary myoblasts and tenoblasts extracted from mice hind limbs are seeded on the threads.

## Material & Methods

### Extrusion

Threads were produced based on protocols developed in [Rieu tb free]. Briefly, collagen solutions (extracted from rat tail tendons) were concentrated at 60 mg/mL in 3 mM HCl 0.8 mM citrate. Fibrinogen solutions were made from fibrinogen stock solutions (Merck) at 40 mg/mL by addition of HCl to reach 60 mM HCl. To obtain fibrin solutions, the acidic fibrinogen solution was supplemented with CaCl_2_ (10 mM final) and thrombin (Sigma, 20 U/mL final). Such solution was either directly mixed with collagen at a ratio 1:1 (volume:volume) to prepare mix solution, or incubated 10 mn to reach suitable viscosity for pure fibrin extrusion.

Solutions were then transferred to a 1 mL syringe and centrifugated for 8 mn at 3000 x g to remove bubbles. They were then extruded in 100 mM HEPES 2.5 w.% PEG solutions, pH 7.4, through a 370 µm needle. The resulting threads (Fig.1.A) were left for at least 10 days in the HEPES/PEG solution and then rinsed at least three times with cell culture medium and left in it for 24 hours to remove PEG.

**Figure 1:**
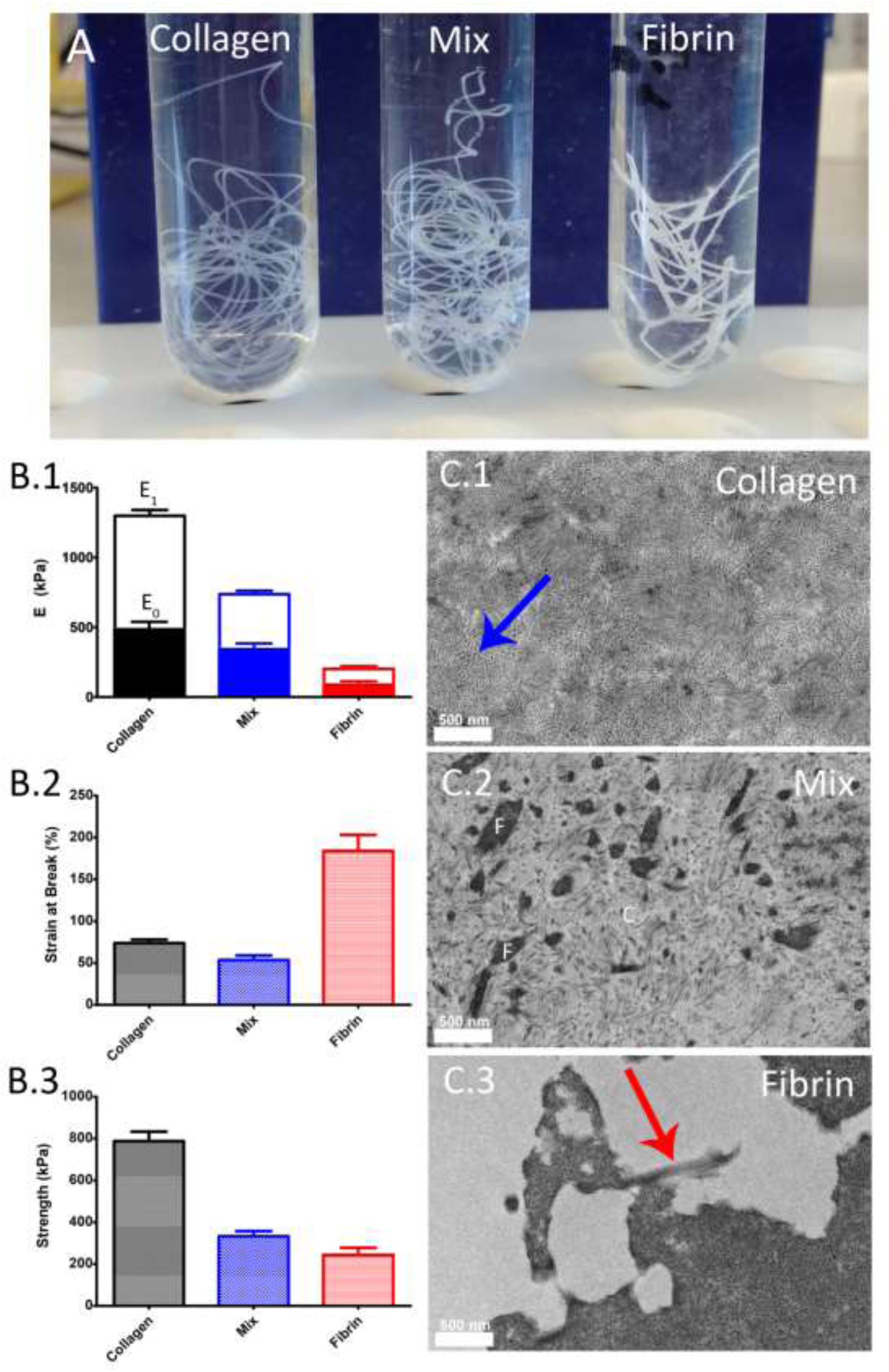
Threads structural & mechanical properties. (A) Macroscopic photos of the three types of threads: collagen, mix and fibrin. (B) Mechanical properties of the three types of threads: (B.1) low-strain E0 (full histogram) and high-strain E1 (empty histogram) moduli, (B.2) strain at break, (B.3) ultimate tensile strength (UTS). Cross sections of the (C.1) collagen, (C.2) mix and (C.3) fibrin threads as observed by transmission electron microscopy. Blue arrow points at collagen fibers aligned along the thread. Red arrow points at the fibrous-like structure at the surface of the fibrin aggregates.

### Tensile Testing

Tensile testing was done on an Instron 3343 machine equipped with a 10N load-cell and a watertight column to perform the tests in hydrated conditions. Tests were performed in PBS 1X, to mimic physiological conditions and thus give an idea of the possible mechanical behavior *in vivo*. Threads were immobilized into clamps with Velcro tape to prevent sliding of the threads. Strain was calculated with the displacement of the clamps.

### Transmission Electron Microcopy

Hydrated samples, crosslinked with PFA, glutaraldehyde and osmium tetroxide 4 wt. %, were subsequently dehydrated using baths with increasing concentrations in ethanol. They were then progressively transferred to propylene oxide and to araldite resin prior to full embedding in pure araldite. After inclusion, 70 nm ultrathin sections (Leica microtome) were made and contrasted with uranyl acetate and observed on a transmission electron microscope FEI Tecnai Spirit G2 operating at 120 kV. Images were recorded on a Gatan Orius CCD camera.

### Cell isolation

Satellite cells were extracted from the hindlimb muscles (hamstring muscle group, quadriceps, tibialis, extensor digitorum longus, gastrocnemius, soleus, gluteus) of 2 to 3 months-old mice while tenoblasts were extracted from the mice Achilles tendons.

Muscles were dissected and minced to a pulp using forceps then digested with collagenase B solution (0.75 U/mL final, from Roche) with dispase II (1.2 U/mL final, from Roche) and CaCl2 (2.5 mM) at 37 °C for 40 minutes. Preparations were washed twice in culture media, filtered through a 70 µm cell strainer and subjected to immunomagnetic cell sorting using a “satellite cell isolation” kit and a MACS “midi” column (Miltenyi Biotech).

Tendons were either digested in 600 U/mL of collagenase II (Worthington) or in 1.5 U/mL of collagenase B (Roche) at 37°C for 18 hours, mechanically destroyed by up and down pipetting and filtered through a 100 µm cell strainer.

Primary myoblasts and tenoblasts were then amplified and cultured on collagen-coated plates for several passages.

### Thread seeding

Threads were cut in approximately 1-cm-long pieces. They were seeded with myoblasts or tenoblasts at 10^5^ cell/cm^2^ in 24-well plates. Cells used were between P5 and P10 for both types. For tenoblasts, 3 threads of each condition were seeded with a first cell line and 1 thread with a different one. For myoblasts, two threads were seeded with a first line and two others with a second one.

Growth medium (high-serum medium - 79 % Hams F-10 from Gibco, 20 % fetal bovine serum, 1 % antibiotic-antimycotic Gibco, with 2.5 ng/mL recombinant human fibroblast growth factor R&D Systems) and fusion medium (low-serum medium - 94 % Dulbecco’s Modified Eagle’s Medium, Gibco, 5 % horse serum, 1 % antibiotic/ antimycotic Gibco) were changed every other day. Unless specified, the differentiation process involves 7 days of proliferation in the growth medium, followed by differentiation in the fusion medium. Aminocaproic acid (6-aminohexanoic acid, Sigma) at a final concentration of 2 mg/mL was added to both media. Recombinant human Wnt7a (Chinese Hamster Ovary cell line-derived, R&D Systems) at a concentration of 50 ng/mL was used for the culture displayed on Supp.Fig.1.

### 2D culture

Culture of myoblasts in 2D was performed in 24 well plates coated with the different materials. Plates were coated with collagen, mix and fibrin with the same solutions as the ones used for extrusion. A thin layer of the solution was deposited, with further addition of growth medium to trigger fibrillation. Matrigel-coated plates, used to promote fusion, were seeded as a reference (BD Biosciences). They were prepared by incubation at 37°C for 30 mn of a thin layer of Matrigel at 1 mg/mL and subsequent drying under the hood. They were then rehydrated using growth medium. Each well was seeded with a density of 2×10^4^ cells/cm^2^, with triplicates per material, and left 1 day in proliferation medium for the myoblasts to attach and then cultured in fusion medium for 6 days. Aminocaproic acid at a final concentration of 2 mg/mL was added to both media.

### Fluorescence microscopy

To be observed the seeded threads were rinsed with PBS 1X and then fixed with 4 % PFA for 15 minutes at room temperature. After a blocking step with BSA 4% for 45 minutes, cells were stained using primary (Myosin heavy chain: primary hybridoma mouse IgG2b; Myogenin: myogenin antibody (5FD) Santa Cruz biology;) and secondary antibodies (Red: Alexa Fluor 546 goat anti-mouse IgG2b, In vitrogen; Magenta: A647 anti-rabbit, Life Technology). Nuclei were stained with Hoechst (Life Technology). Observations were made on a calibrated EvosR fluorescence inverted microscope.

To assess homogeneity of mix threads, fibrinogen-FITC was used. To label fibrinogen, 20 mg of FITC CeliteR was dissolved in 3 mL of 0.13 M carbonate buffer pH 9.2. Fibrinogen stock solution at 40 mg/mL was then mixed to the FITC solution at a ratio 1:3. The resulting solution was then dialysed 48 hours against 2 L of a solution of 20 mM citrate pH 7.4, with regular changes of the dialysing bath. Final solution was aliquoted and stored at -80°C. For extrusion, 10% in volume of FITC-labelled fibrinogen [Fbg-FITC] was added to fibrinogen stock solution, before addition of HCl, and following the usual protocol afterwards.

### Statistical analysis

For statistical analysis, imaging of every part of the threads was carried out at 4x magnification to minimize deviation due to image sampling. ImageJ software was used to measure cell density. An area taking only the central part of the thread was outlined and measured using ImageJ. The central area was chosen to be approximately two thirds of the thread width. This was done for two reasons. First, the projected area measured on the picture in 2D was smaller than the actual area in 3D, due to the curvature of the threads. For a ratio r = 2/3, the relative error made is around 8% which is reasonable. Second, due to the curvature of the thread, the cells outside the central part tend to appear as cell aggregates and are more difficult to distinguish and count. Cells inside the outlined area were manually counted using the cell counter plug-in from ImageJ. To count cells positive to MF20, a composite picture was used with both the blue channel (Hoechst, nuclei) and the red channel (MF20, heavy chain myosin).

Prism software was used to draw the whisker boxes as well as to perform Mann Whitney tests to compare two distributions.

## Results

### Mechanical properties

Results of mechanical tests are displayed on Fig.1.B. First, we noticed that the three types of threads exhibit a strain stiffening behavior. Two Young moduli were then measured: at low strain, noted E_0_, and during strain-stiffening, written E_1_ (Fig.1.B.1). For both moduli, the difference between collagen and fibrin stands out: collagen is much stiffer (E_0_ = 0.57 ± 0.05 MPa, E_1_ = 1.11 ± 0.13 MPa) than fibrin (E_0_ = 0.10 ± 0.02 MPa, E_1_ = 0.21 ± 0.06 MPa). Mix threads exhibit intermediate stiffness (E_0_ = 0.33 ± 0.05 MPa, E_1_ = 0.83 ± 0.19 MPa) between collagen and fibrin.

Fibrin threads exhibit higher strain at break than collagen (251 ± 63 % *vs.* 75 ± 15 %) (Fig.1.B.2) but lower ultimate tensile strength (0.33 ± 0.04 MPa vs. 0.70 ± 0.11 MPa) (Fig.1.B.3). Interestingly, mix threads seem to break at even lower strain than collagen (57 ± 21 %) with an ultimate tensile strength slightly higher than for fibrin (0.42± 0.2 MPa).

### Structure

Investigation of the microstructure of threads was performed on ultrathin transverse sections of threads using Transmission Electron Microscopy (Fig.1.C). Collagen threads are made of thin fibers with different orientations, among which a significant part is pointing toward the direction of observation, *i.e.* along the thread axis (blue arrow, Fig.1.C.1). Fibrin threads exhibit dense aggregate-like structures with some fibrous-like structures on the outside (red arrow, Fig.1.C.3). Mix threads structure is composed of small islands of fibrin materials (noted F on Fig.1.C.2), embedded into a collagen matrix (C Fig.1.C.2), made of thicker fibers than the ones observed in collagen alone.

### Tenoblasts culture on threads

Tenoblasts were seeded on the three types of threads and observed after 4 days in high-serum medium (Fig.2.A) and after 7 days in high-serum medium plus 3 days in low-serum medium (Fig.2.B). After 4 days, we observed that tenoblasts on collagen threads would form cell “crusts” that detached from the thread (Fig.2.A.1). On mix and fibrin threads, tenoblasts cover threads with high cell density, without cell crust detaching from them (Fig.2.A.2 & A.3 resp.). The detachment of cells causes the cell density to drop on collagen (median ≈ 200 cell/mm^2^) compared to mix (med. ≈ 1100 cell/mm^2^, p=0.012) and fibrin (med. ≈ 1000 cell/mm^2^, p=0.006) threads, with no significant difference between mix and fibrin (Fig.2.A.4).

**Figure 2:**
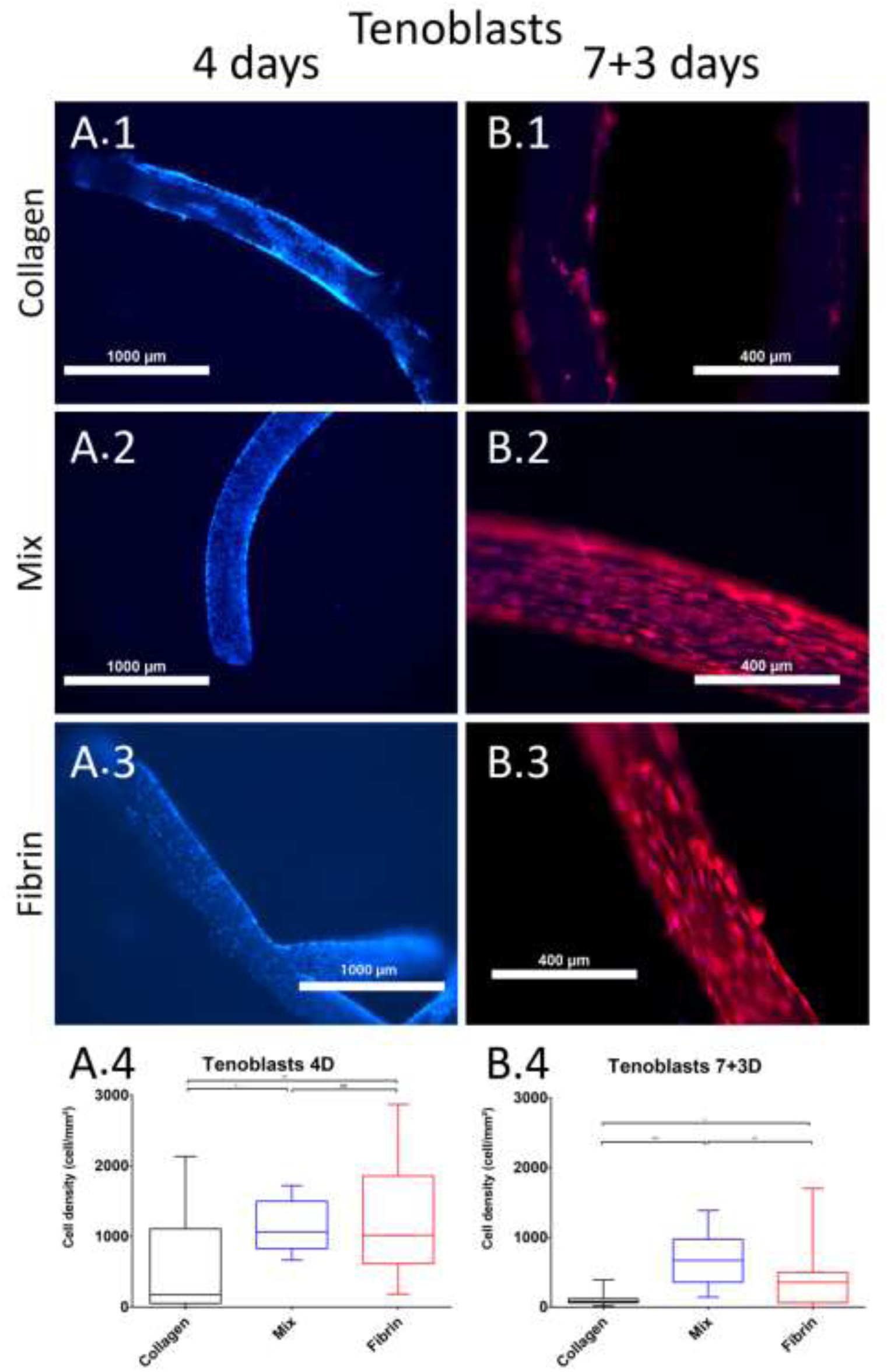
Tenoblast colonization of the three types of threads after day 4 (A) and 7+3 (B). Fluorescence microscope images of a collagen, mix and fibrin threads after 4 days in high-serum medium (A.1, A.2 and A.3 resp) and after 7 days in high-serum medium and 3 days in low serum medium (B.1, B.2 and B.3 resp.), and the resulting tenoblasts densities at day 4 (A.4) and day 7+3 (B.4). Blue staining: Hoechst; red: F-actin.

After 7 days in high-serum medium plus 3 days in low-serum medium, few cells remain on collagen threads (Fig.2.B.1) while mix and fibrin threads still appear covered with cells (Fig.2.B.2 & B.3 resp.). On both types of threads, cells seem to align along the main axis of the threads. Cell density is extremely low on collagen threads (med. < 100 cell/mm^2^), much below the ones on mix threads (med. ≈ 700 cell/mm^2^, p<0.0001) and fibrin threads (med. ≈ 350 cell/mm^2^, p=0.020) (Fig.2.B.4). Compared to cell density after 4 days in high-serum medium, a decrease is observed on mix threads (p=0.0014) and even more on fibrin threads (p=0.0016), resulting in a lower density on fibrin than on mix threads (p=0.0059). The decrease occurs during the low-serum culture, as no significant change in cell density could be observed for both types of threads at day 7 (data not shown).

### Myoblasts culture on threads

Seeding of myoblasts on the three types of threads resulted in a poor adhesion of the cells on collagen threads after 4 days in high-serum medium (Fig.3.A.1) while mix and fibrin threads were well colonized (Fig.A.2 & A.3 resp.). Same trend was observed after 3 days of differentiation, with rare cells found on collagen (Fig.3.B.1) and well colonized mix and fibrin threads (Fig.3.B.2 & B.3 resp.). Most myoblasts on mix and fibrin threads were positive to MF20 staining exhibiting production of myosin heavy chains, a late-differentiation marker. Quantitatively, after 4 days, cell density was dramatically lower on collagen (med. = 3 cell/mm^2^) than on mix (med. ≈70 cell/mm^2^, p<0.001) and fibrin threads (med. ≈ 90 cell/mm^2^, p<0.001) (Fig.3.A.4). After 3 days of differentiation, cellular densities exhibited higher median value on both mix (med. ≈200 cell/mm^2^ p=0.035) and fibrin (med. ≈ 300 cell/mm^2^, ns) compared to 4 days of proliferation. Myoblasts on collagen were too scarce to enable statistically relevant count. No significant difference in cell density between mix and fibrin could be observed either after 4 days of proliferation or 3 days in differentiation. A slight difference in MF20 positive myoblast ratio was however noticed with 84 % of MF20+ cells on mix compared to the 80% on threads (p=0.015) (day 7+3). Interestingly, myoblasts remained round-shaped on both types of threads and no trace of spindle-shaped or fused cells could be observed. Prolonged culture in low-serum medium (5 days) and in presence of Wnt7a at 50 ng/mL fostered faint spindle shape but no fusion as seen on Supp. Fig. 1.A &B. As a reference, same line cultured on a 2D substrate coated with Matrigel exhibited high number of myotubes (Supp. Fig. 1.C), in spite of the same cellular density (Supp. Fig. 1.D).

**Figure 3:**
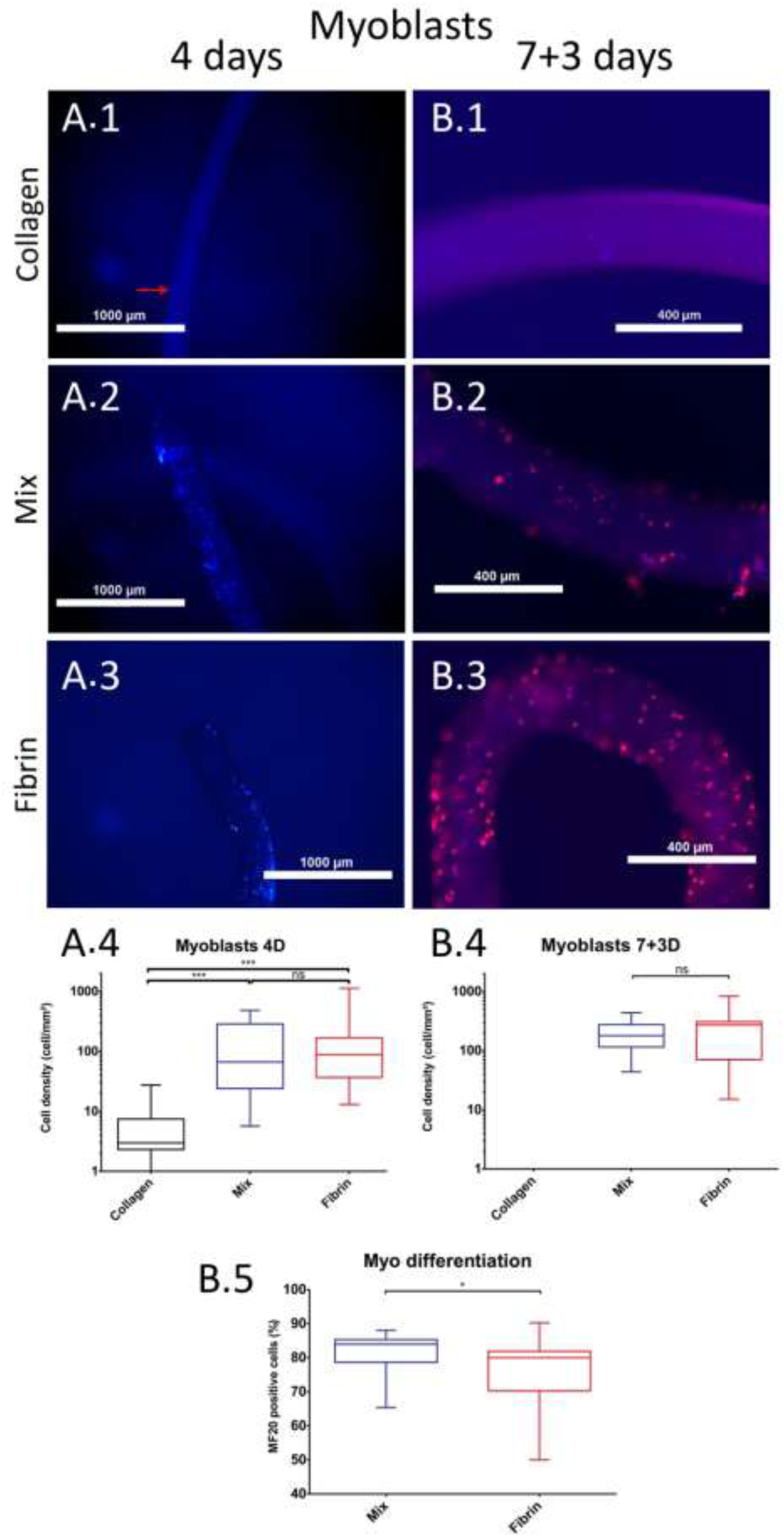
Myoblast colonization of the three types of threads after day 4 (A) and 7+3 (B). Fluorescence microscope images of a collagen, mix and fibrin threads after 4 days in high-serum medium (A.1, A.2 and A.3 resp.) and after 7 days in high-serum medium plus 3 days in low serum medium (B.1, B.2 and B.3 resp.), and the resulting myoblasts densities at day 4 (A.4) and day 7+3 (B.4). (B.5) Proportion of MF20 positive myoblasts over the whole number of myoblasts on mix and collagen threads. Blue staining: Hoechst; red: F-actin. Staining Blue: Hoechst, Red: myosin heavy chains MF20.

**Supp. Fig. 1:**
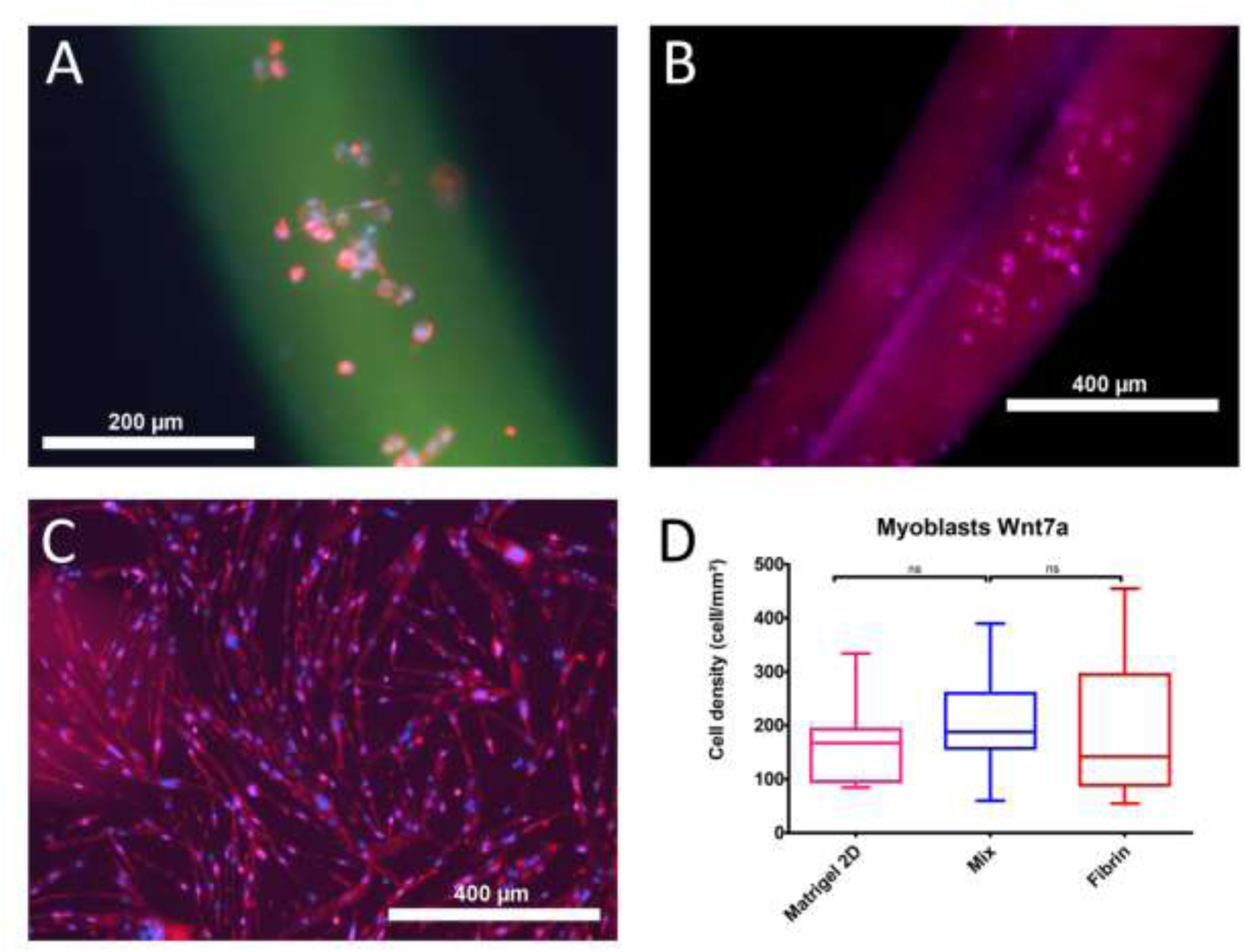
Primary myoblasts cultured for 7 days in proliferation and 5 days in differentiation with Wnt7a at 50 ng/mL. Immunofluorescence staining images of myoblasts on mix (A) and fibrin (B) threads and on Matrigel^®^-coated plates (C) (blue: Hoechst, red: myosin heavy chains, magenta: myogenin, green: fibrin marked FITC). (D) Resulting cell densities on plates and threads.

### Myoblast culture in 2D

To investigate the role played by the threads substrates on myoblasts fusion, compared to 2D substrates, the same solutions as the ones used for thread production were coated on well-plates. Myoblasts were also seeded on Matrigel-coated wells as a control. As seen on Fig.4, several features stand out. Myoblasts seeded on collagen exhibited scarce spindle shapes and extremely low numbers of fused cells (Fig.4.A). On the contrary, myoblasts cultured on mix (Fig.4.B) and fibrin wells (Fig.4.C) could elongate and formed myotubes, with an apparent number of myotubes being higher on mix. Myoblasts on Matrigel formed many more and much larger myotubes (Fig.4.C).

**Figure 4:**
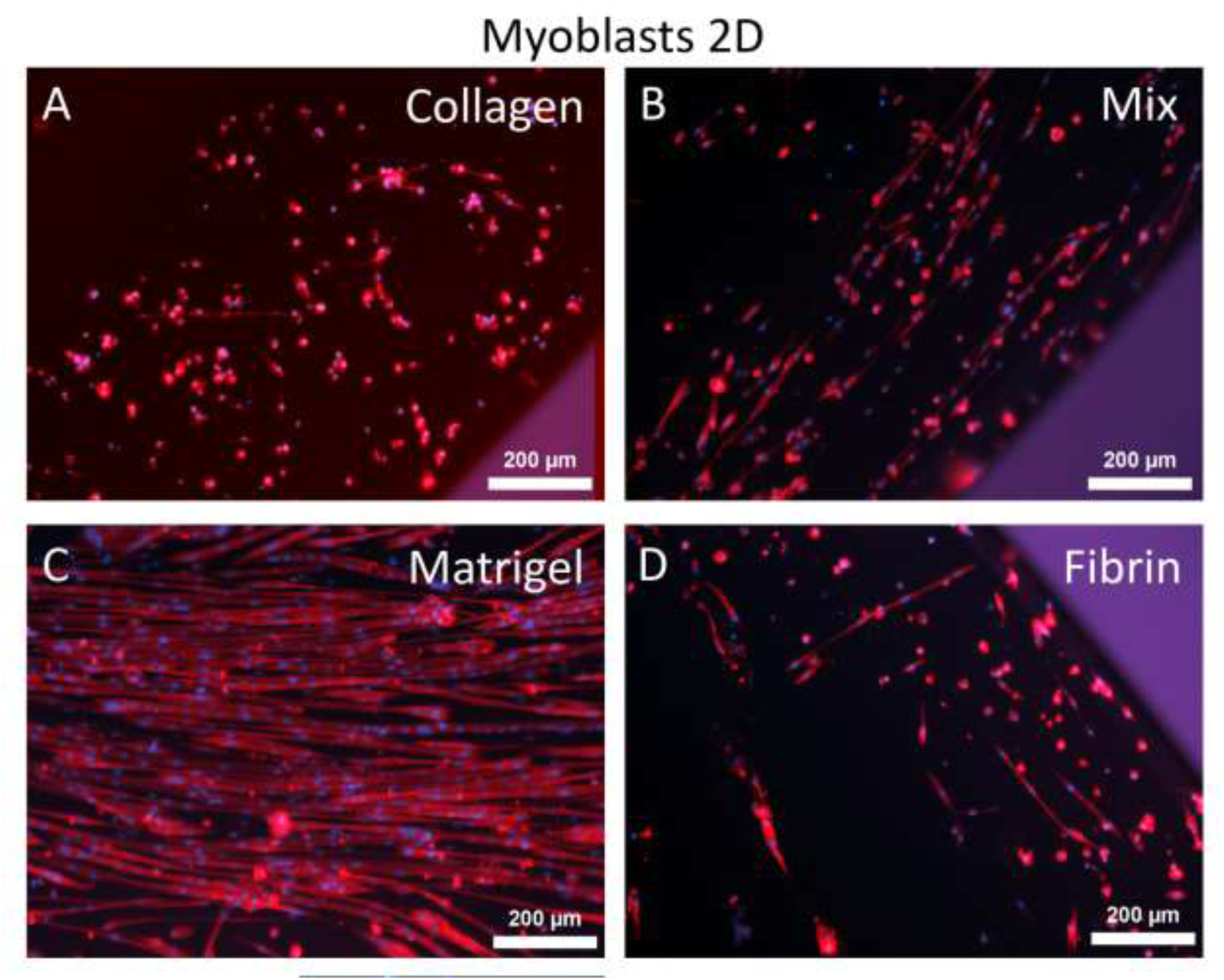
Primary myoblasts cultured for 1 day in proliferation and 6 days in differentiation. Immunofluorescence staining images of myoblasts on collagen (A), mix (B), Matrigel^®^ (C) and fibrin (D) coated plates (blue: Hoechst, red: myosin heavy chains

## Discussion

We have produced in this work three types of threads, one made of pure collagen, one of pure fibrin and the last one of a mix of collagen and fibrin at a ratio 3:2 in mass. These threads differing in terms of present also different mechanical properties.

### Mechanical properties

In tensile testing, we noticed that the three types of threads exhibit a strain stiffening behavior characteristic of fibrin and collagen materials ^46^. As expected, collagen threads exhibited stiffer behavior, with higher strength, and lower strain at break compared to fibrin, which could be stretched up to approximately 250%. In accordance with first intuitions, mix threads had intermediate stiffness and strength between fibrin and collagen. Mix threads had however lower strain at break than collagen.

Interestingly, our results are consistent with the ones obtained by Lai *et al.* ^47^ who studied mechanical properties of lower concentration (2 to 3 mg/mL in proteins) collagen/fibrin gels with different proportions from 100 % collagen to 100 % fibrin to understand the nature of the networks and their relationships in fibrin/collagen composites. The reported values of modulus and strain at failure of composites at 60 % collagen are closer to those for pure collagen than for pure fibrin. They demonstrated that such behavior could not be explained by a simple parallel (*i.e.* non interacting networks) or series (*i.e.* co-dependent networks) model of collagen and fibrin materials. They partially explained mechanical properties obtained in this study in a second article using a model of non-percolating network in which islands of the diluted protein are embedded in the concentrated one ^48^. Such conclusion is in total agreement with the structure observed in mix threads, with islands of fibrin inside the collagen matrix (Fig.1.C.2). While mix threads are macroscopically homogeneous as observed on threads made from FITC-labelled fibrinogen, it seems that a phase separation occurs microscopically, resulting in islands of fibrin in a collagen matrix, for the ratio collagen:fibrin used here.

Structure of collagen threads with small fibrils with local orientation is typical of highly concentrated collagen scaffolds ^49^ and was already observed in threads extruded in high osmotic pressure buffer ^50^. Fibrin’s structure is similar to the structure we observed in fibrin threads extruded in Dulbecco’s Modified Eagle Medium with a fibrillar structure on the outside of the aggregates instead of the thin bridging fibers, probably due to high osmotic pressure exerted by the high PEG concentration in the extrusion buffer. Same structure is observed for fibrin islands in mix threads. The small size of islands and the low distance between them (<1µm) ensure that cells can access both collagen and fibrin materials. It may be interesting to think about the “clustering” effect of these islands on cellular response, compared to a situation where collagen and fibrin would be thouroughly mixed, as clustering effect has shown to have paramount effect on cellular response.

### Tenoblasts

Several features stood out from the culture of tenoblasts on collagen, mix and fibrin threads. First, tenoblasts rapidly proliferated rapidly on collagen threads, due to high affinity, and rapidly formed a crust around the threads, made of interconnected tenoblasts and secreted ECM ^51,52^. After only 4 days in culture the crust detached from collagen threads suggesting that collagen is not a suitable substrate to enable long-term culture of tenoblasts. The difference in mechanical properties as well as the tendency of cells to contract their surrounding matrix may cause the observed crust-detaching effect.

Second, the cell density drops on mix and especially on fibrin threads between day 7 and day 7+3. This drop is likely to be caused by the switch from high-serum to low-serum medium. While mix and fibrin had similar densities at day 4 and 7, cell density is lower at day 7+3 on fibrin than on mix. The combination of collagen and fibrin thus seems beneficial to maintain tenoblasts survival in low serum.

On fibrin threads, cell density seems rather stable during the 7 days of proliferation. After 7 days (data not shown), some small cell plaques seem to detach but without significantly affecting cell density when compared to density at day 4. After 3 days in differentiation medium, cell density decreased. This may be due to the change to low serum-content medium, or to the detachment due to confluence in a similar fashion as for collagen.

Third, tenoblasts seem to spontaneously align along the direction of the thread on both mix and fibrin threads. This may be explained by a minimization of the stress on internal structure due to bending as already pointed out ^53^.

### Myoblasts

The observed low adherence of myoblasts on collagen threads may appear surprising as primary myoblasts are routinely cultured on collagen-coated plates for proliferation ^54^. Actually, on collagen-coated dishes myoblasts adopt a round shape demonstrating low affinity to collagen whereas they strongly adhere on coating containing laminin and/or fibronectin such as Matrigel^®^ and adopt tapered shape ^55^. In fact, myoblasts do not bind directly to collagen but through fibronectin ^30^, and the one provided by the addition of serum is supposed to be sufficient to permit adhesion of myoblasts ^56^.

Given the higher densities on mix and fibrin threads, myoblasts seemed to adhere more on fibrin. Myoblasts can bind directly to fibrin through integrin receptors ^57^. The cell density on mix threads was each time comparable to the one on fibrin. It is surprising that mix threads induced similar response as pure fibrin, given the relatively low content in fibrin compared to collagen. Fibrin islands contained in mix threads were thus sufficient to provide similar adhesion of cells to threads as on pure fibrin. Given the absence of adhesion of myoblasts on collagen threads, we can hypothesize that myoblasts attach to fibrin islands instead of collagen matrix, and thus “feel” a fibrin composition. However, given the small size of islands embedded in the collagen matrix, it is probable that cells, when pulling on focal adhesion, rather “feel” the stiffness of the collagen matrix. Mix materials may then appear as a way to decouple the composition and mechanical properties of the substrate and thus show that the lack of adherence of myoblasts to collagen threads is rather due to the poor binding to collagen compared to fibrin, rather than a too high stiffness of the substrate.

Concentration of proteins in threads may also play a role. Gerard *et al.* ^58^ reported that concentration of fibrin had an effect on myoblasts entrapped in a fibrin gel. They demonstrated that loose gels (3mg/mL) promoted higher proliferation and metabolic activity compared to 50 mg/mL gels.

The absence of fusion of myoblasts into myotubes was unexpected as fibrin has been widely used for muscle constructs ^43–45,59^ and is believed to be the gold-standard material ^60^. However, myoblasts on threads expressed myogenin and heavy myosin chains, two markers of muscle cell terminal differentiation. The high density of multinucleated myotubes on 2D Matrigel control proves that the absence of fusion on the threads is not due to low cellular density or to an inability of the primary myoblasts we used to fuse. We tested on differentiated cells on threads the stimulation by Wnt7a that, as previously reported, is an inducer of muscle cell fusion ^61^. We only observed a modest effect on cell sprawling and polarization ^62^, still too low to enable cell fusion. Given the fact that most cells expressed were terminally differentiated, and kept a round “myoblastic” shape on threads, it seemed that the absence of fusion is due to a problem of adhesion of cells to the matrix.

The comparison between culture on threads and 2D culture on coated wells enables to point out two distinct reasons that seem to play a role in the absence of fusion of myoblasts. The first is due to the composition of the thread. Collagen alone seems to be particularly non-permissive to myocyte fusion. Differentiation of myoblasts without fusion has already been observed on collagen-coated plates ^63^. On the contrary to collagen, fibrin-coated plates comports multiple myotubes. The combination of collagen and fibrin in mix plates may have a synergetic effect and as they seem to promote even more myotube fusion. Number and size of myotubes are however much lower than on Matrigel, which contains multiple ECM components. The lack of fusion may thus result from the lack of certain components of the ECM such as laminin, shown to be paramount for cell fusion ^64^ and often used to promote myoblasts adhesion and fusion ^65,66^.

A clear difference could however be observed between myoblasts seeded on 2D fibrin or mix, compared to myoblasts seeded on threads. Multiple myotubes and spindle-shaped cells could be observed in 2D while no fusion was witnessed on threads, and cell polarization and elongation was tediously and limitedly obtained using Wnt7a. As the materials used in 2D and threads are identical, the absence of fusion is likely to be due to the topology of the threads, *ie.* the curvature. Curvature has already been reported to influence cell migration and differentiation ^67^ but to our knowledge, no study has shown that substrate curvature inhibited myoblast fusion. In 1994, Rovensky *et al.* reported that cells with straight actin filament such as fibroblasts tend to align in the direction of lower curvature to minimize stress ^53^. We can hypothesize that the myoblasts do not fuse and do not even elongate because they are highly sensitive to internal stress.

It is known that muscle satellite cells do not fuse on an isolated myofibre while they fuse once plated on a Matrigel^®^-coated plate ^68^. The shape and curvature of the substrate may then inhibit fusion. Myoblasts on our threads seemed to behave in the same fashion as satellite cells on isolated myofibres, adopting the same shape and unable to fuse. The topological inhibition is surprising given the difference of dimension compared to myofibers with diameters of a few tens of micrometers ^69,70^. The curvature inhibition may prevent differentiation of satellite cells at the surface of undamaged myofibers.

The threads developed may then represent an interesting model of isolated myofibers. This is also encouraging for muscular reconstruction as satellite cells adopt such a behavior when anchored to an isolated fiber but behave differently in a 3D environment, in the niche comprised between multiple fibers. Use of multiple braided threads for instance, possibly held together by a loose gel, may then form a niche model for myoblasts and effectively promote myotube formation and muscle reconstruction.

## Conclusion

We developed here collagen, fibrin and mixed threads and used them as substrates of different mechanical properties, collagen threads being stiffer than fibrin ones. Mix threads demonstrated properties similar to collagen due to their non percolating fibrin island network in the collagen matrix. Such substrates enable to decouple mechanical properties from chemical interaction by providing fibrin islands for the cells to adhere with the stiffness of the embedding collagen network. Both primary tenoblasts and myoblasts on mix threads behave similarly to fibrin threads, rather than collagen, highlighting the importance of the chemical nature of the cell/matrix interaction.

Even though myoblasts differentiated on mix and fibrin threads, they did not fuse. Investigations carried out seem to point out a possible inhibition of fusion due to substrate curvature. Further characterization of such phenomenon should be undertaken as it may provide a better understanding of the satellite cells differentiation commitment all the more since prevention of satellite cell differentiation is a challenge for muscular reconstruction ^71^. Indeed, surface curvature seems to be a parameter of choice to foster or not the differentiation of satellite cells and ensure differentiation only upon fiber damage.

The work reported here is a first step toward myo-tendinous junction modelisation. The next step would be to co-culture both tenoblasts and myoblasts on our substrates, as we demonstrated their viability in the same medium, to understand the cooperative role they may play. As fibrin, collagen and mix threads are produced in the same conditions, collagen:fibrin ratio of mix threads can be tuned to investigate cell response to substrate and find the optimal composition for tenoblasts and myoblasts. Even further, pluripotent stem cells could be used and composition of mix threads adapted to favor tenogenic or myogenic commitment.

## Notes

### Competing Interest Statement

The authors have declared no competing interest.

